# ReaDDy 2: Fast and flexible software framework for interacting-particle reaction dynamics

**DOI:** 10.1101/374942

**Authors:** Moritz Hoffmann, Christoph Fröhner, Frank Noé

## Abstract

Interacting-particle reaction dynamics (iPRD) combines the simulation of dynamical trajectories of interacting particles as in molecular dynamics (MD) simulations with reaction kinetics, in which particles appear, disappear, or change their type and interactions based on a set of reaction rules. This combination facilitates the simulation of reaction kinetics in crowded environments, involving complex molecular geometries such as polymers, and employing complex reaction mechanisms such as breaking and fusion of polymers. iPRD simulations are ideal to simulate the detailed spatiotemporal reaction mechanism in complex and dense environments, such as in signalling processes at cellular membranes, or in nano- to microscale chemical reactors. Here we introduce the iPRD software ReaDDy 2, which provides a Python interface in which the simulation environment, particle interactions and reaction rules can be conveniently defined and the simulation can be run, stored and analyzed. A C++ interface is available to enable deeper and more flexible interactions with the framework. The main computational work of ReaDDy 2 is done in hardware-specific simulation kernels. While the version introduced here provides single- and multi-threading CPU kernels, the architecture is ready to implement GPU and multi-node kernels. We demonstrate the efficiency and validity of ReaDDy 2 using several benchmark examples. ReaDDy 2 is available at the https://readdy.github.io/ website.

## 1 Introduction

The physiological response of biological cells to stimuli can be a many-stage process. A widely studied example is the MAPK pathway [56, 45]. Many of such signaling pathways incorporate G-protein coupled receptors (GPCR) [48] and cyclic adenosine monophosphate (cAMP) [8]. These are related to various diseases [47, 17, 1]. An extracellular stimulus can activate the membrane bound GPCRs and lead to localized synthesis of cAMP as second messengers. Their transport through the cell is diffusive, however due to the geometry of cellular compartments cAMP molecules are non-uniformly distributed [2, 27]. Their presence needs to be resolved in space and time to understand their function.

Particle-based reaction dynamics (PBRD) simulations [19, 52, 16] are amongst the most detailed approaches to model reaction kinetics computationally as they simulate each reactive molecule as a particle and therefore can be used as a tool to investigate systems of the aforementioned kind. Reactions can occur when reactive particles are in proximity, resembling the physical process. PBRD is suitable when the spatial distribution of molecules does not equilibrate rapidly and must therefore be resolved, and some reactants are locally scarce, such that their discrete number must be kept track of [9]. There is a wide range of simulation tools for PBRD [43], including Smoldyn [4], MCell [31], Cell++ [37], eGFRD [45], mesoRD [24], spatiocyte [5], SpringSaLaD [33], and SRSim [22].

PBRD simulations usually contain purely reactive particles that are not subject to interaction forces, e.g., to model space exclusion with repulsive interactions or clustering with attractive interactions. On the other hand, molecular dynamics (MD) simulations are designed to model particle dynamics including complex interactions between the particles or particles and an external field. The particles in MD simulations are often atoms or groups of atoms and higher-order structures such as molecules are represented by topology graphs that define the bonding structure between particles and thus, together with a MD force field, imply which pair, triplets and quadruplets or particles interact by means of bond, angle and torsion potentials. While reactive force fields [26, 30, 50] include reactivity on the chemistry scale, and soft matter MD simulation tools include breakable bonds [6, 32], current MD models and simulation packages do not incorporate generic particle reactions.

Interacting-particle reaction dynamics (iPRD) was introduced in [42] to combine the benefits of PBRD and MD simulations by modeling particle-based reaction dynamics while enabling full-blown interactions between particles as well as particles and the environment. On mesoscopic length scales these interactions can be used to induce structure, e.g., volume-exclusion in crowded systems [25, 42], clustering of weakly interacting macromolecules [49], restriction of diffusing particles to arbitrarily-shaped membranes [42, 23, 43]. Furthermore it allows to study the large-scale structure of polymers and membranes [36]. When not only considering interactions but also reactions, a wide range of reactive biochemical systems are in the scope of the model. For example, the reaction dynamics of photoreceptor proteins in crowded membranes [39] including cooperative effects of transmembrane protein oligomers [23] have been investigated. Another example is endocytosis, in which different proteins interact in very specific geometries [35, 40].

The price of resolving these details is that the computation is dominated by computing particle-particle interaction forces. Although non-interacting particles can be propagated quickly by exploiting solutions of the diffusion equation [38, 51, 52, 15], interacting particles are propagated with small time-steps [53, 54], restricting the accessible simulation timescales whenever parts of the system are dense. As this computational expense is not entirely avoidable when the particle interactions present in iPRD are needed to model the process of interest realistically, it is important to have a simulation package that can fully exploit the computational resources.

Here we introduce the iPRD simulation framework ReaDDy 2, which is significantly faster, more flexible, and more conveniently usable than its predecessor ReaDDy 1 [42, 10]. Specifically, ReaDDy 2 includes the following new features:

### • Computational efficiency and flexibility

ReaDDy 2 defines computing kernels which perform the computationally most costly operations and are optimized for a given computing environment. The current version provides a single-CPU kernel that is four to ten times (depending on system size) faster than ReaDDy 1, and a multi-CPU kernel that scales with 80% efficiency to number of physical CPU cores for large particle systems (Sec. 3.2). Kernels for GPUs or parallel multi-node kernels can be readily implemented with relatively little additional programming work (Sec. 3).

### • Python user interface

ReaDDy 2 can be installed via the conda package manager and used as a regular python package. The python interface provides the user with functionality to compose the simulation system, define particle interactions, reactions and parameters, as well as run, store and analyze simulations.

### • C++ user interface

ReaDDy 2 is mainly implemented in C++. Developers interested in extending the functionality of ReaDDy 2 in a way that interferes with the compute kernels, e.g., by adding new particle dynamics or reaction schemes, can do that via the C++ user interface.

### • Reversible reaction dynamics

ReaDDy 2 can treat reversible iPRD reactions by using steps that obey detailed balance, as described in [21] (iPRD-DB), and thus ensure correct thermodynamic behavior for such reactions (Sec. 4.1).

### • Topologies

We enable building complex multi-particle structures, such as polymers, by defining topology graphs (briefly: topologies, see Sec. 2.4). As in MD simulations, topologies are an efficient way to encode which bonded interactions (bond, angle and torsion terms) should act between groups of particles in the same topology. Note that particles in topologies can still be reactive. For example, it is possible to define reactions that involve breaking or fusing polymers (Sec. 4.4).

This paper summarizes the features of ReaDDy 2 and the demonstrates its efficiency and validity of ReaDDy 2 using several benchmarks and reactive particle systems. With few exceptions, we limit our description to the general features that are not likely to become outdated in future versions. Please see https://readdy.github.io/ for more details, tutorials and sample code.

## 2 interacting-Particle Reaction Dynamics (iPRD)

The ReaDDy 2 simulation system consists of particles interacting by potentials and reactions (Fig. 1) at a temperature *T*. Such a simulation system is confined to a box with repulsive or periodic boundaries. A boundary always has to be either periodic or be equipped with repulsive walls so that particles cannot diffuse away arbitrarily. To simulate iPRD dynamics in complex architectures, such as cellular membrane environments with specific shapes, additional potentials can be defined that confine the particle to a sub-volume of the simulation box (Sec. 2.2).

**Figure 1:**
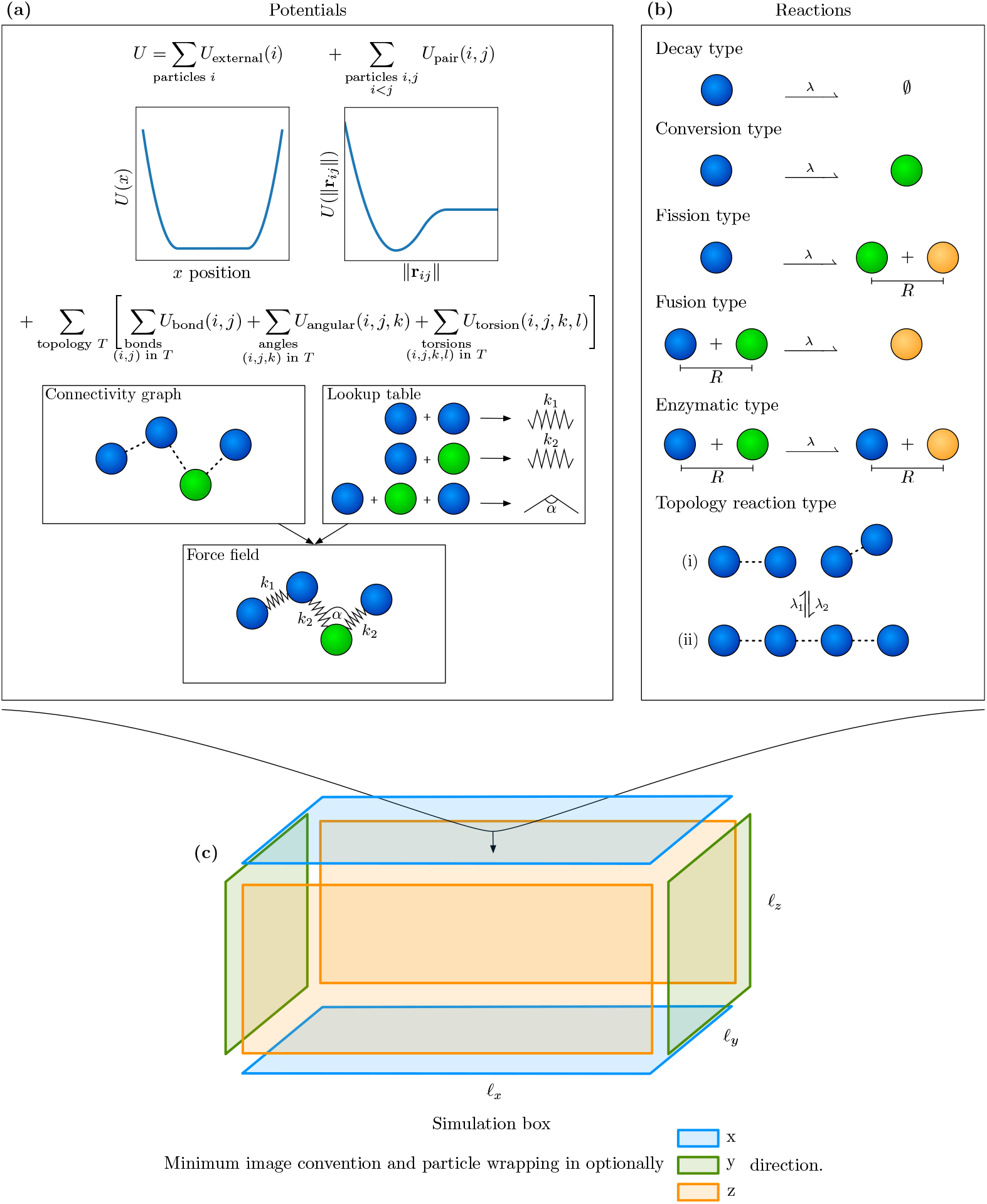
ReaDDy 2 simulation model. **(a)** Potentials: Particles are subject to position-dependent external potentials, such as boundary potentials or external fields and interaction potentials involving two, three or four particles. As in molecular mechanics force fields, bonded potentials are defined within particle groups called “topologies” whose bonding structure is defined by a connectivity graph. **(b)** Reactions: Most reactions are unimolecular or bimolecular particle reactions. Topology reactions act on the connectivity graphs and particle types and therefore change the particle bonding structure.

### 2.1 Interacting particle dynamics

ReaDDy 2 provides a developer interface to flexibly design models of how particle dynamics are propagated in time. The default model, however, is overdamped Langevin dynamics with isotropic diffusion as this is the most commonly used PBRD and iPRD model. In these dynamics a particle *i* moves according to the stochastic differential equation:

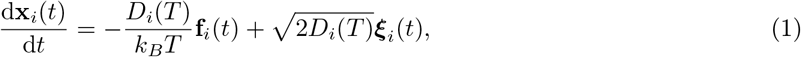

where **x**_*i*_(*t*) ∈ ℝ^3^ contains the particle position at time *t*, *D_i_*(*T*) is its diffusion coefficient, *k_B_* is the Boltzmann constant, and *T* the system temperature. The particle moves according to the deterministic force **f**_*i*_ and the stochastic velocity 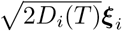 in which ***ξ***_*i*_ are independent, Gaussian distributed random variables with moments

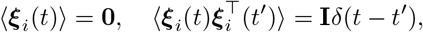

where **I** is the identity matrix. The stochastic terms ***ξ***_*i*_ and ***ξ***_*j*_ are uncorrelated for particles *i* ≠ *j*. In ReaDDy 2 the default assumption is that the diffusion coefficients *D_i_*(*T*) are given for the simulation temperature *T*. Additionally, we offer the option to define diffusion coefficients for a reference temperature *T*_0_ = 293*K* and then generate the diffusion coefficients at the simulation temperature *T* by employing the Einstein-Smoluchowski model for particle diffusion in liquids [55, 18]:

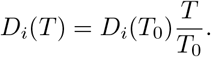

This way, simulations at different temperatures are conventient while only having to specify one diffusion constant. Using this model,the dynamics are

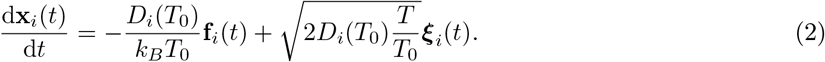

This means that the mobility is preserved if the temperature changes and (1) is recovered for *T* = *T*_0_.

The simplest integration scheme for (1), (2) is Euler-Maruyama, according to which the particle positions evolve as:

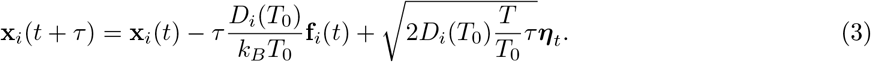

Where *τ >* 0 is a finite time step size and ***η***_*t*_ ~ 𝒩 (0, 1) is a normally-distributed random variable. The diffusion constant *D_i_* effects the magnitude of the random displacement. The particles’ positions are loosely bound to a cuboid simulation box with edge lengths *ℓ_x_, ℓ_y_, ℓ_z_* (Fig. 1). If a boundary is non-periodic it is equipped with a repulsive wall given by the potential

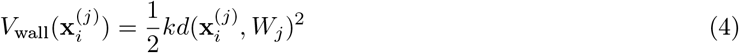

acting on every component *j* of the single particle position **x**_*i*_, where *k* is the force constant, 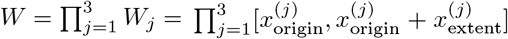 the cuboid in which there is no repulsion contribution of the potential, and *d*( *, W_j_*) := inf *d*( *, w*): *w* ∈ *W_j_* the shortest distance to the set *W_j_*. The cuboid can be larger than the simulation box in the periodic directions. In non-periodic directions there must be at least one repulsive wall for which this is not the case.

Due to the soft nature of the walls particles still can leave the simulation box in non-periodic directions. In that case they are no longer subject to pairwise interactions and bimolecular reactions however still are subject to the force of the wall pulling them back into the box.

Other types of dynamical models and other integration schemes can be implemented in ReaDDy 2 via its C++ interface. For example, non-overdamped dynamics, anisotropic diffusion [54, 34], hydrodynamic interactions [20] or employing the MD-GFRD scheme to make large steps for noninteracting particles will all affect the dynamical model and can be realized by writing suitable plugins.

### 2.2 Potentials

The deterministic forces are given by the gradient of a many-body potential energy *U* (Fig. 1a):

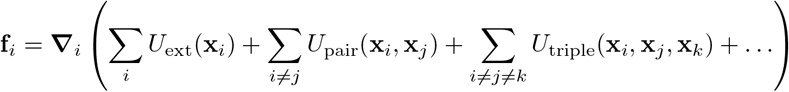

The potentials are defined by the user. ReaDDy 2 provides a selection of standard potential terms, additional custom potentials can be defined via the C++ interface and then included into a Python simulation script.

External potentials only depend on the absolute position of each particle. They can be used, e.g., to form softly repulsive walls (4) and spheres, or to attach particles to a surface, for example to model membrane proteins. Furthermore the standard potential terms enable the user to simulate particles inside spheres and exclude particles from a spherical volume. The mentioned potential terms can also be combined to achieve more complex geometrical structures. Pair potentials generally depend on the particle distance and can be used, e.g., to model space exclusion at short distances, or to model screened electrostatic interactions between particles.

ReaDDy 2 has a special way of treating interaction potentials between bonded particles. Topologies define graphs of particles that are bonded and imply which particle pairs interact via bond constraints, which triples interact via angle constraints, and which quadruplets interact via a torsions potential. See Sec. 2.4 for details.

### 2.3 Reactions

Reactions are discrete events, that can change particle types, add, and remove particles (Fig. 1b). Each reaction is associated with a microscopic rate constant *λ >* 0 which has units of inverse time and represents the probability per unit time of the reaction occurring. The integration time-steps used in ReaDDy 2 should be significantly smaller than the inverse of the largest reaction rate, and we therefore compute discrete reaction probabilities by:

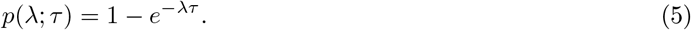

In the software it is checked whether the time step *τ* is smaller than the inverse reaction rate up to a threshold factor of 10, otherwise a warning is displayed as discretization errors might become too large. In general, ReaDDy 2 reactions involve either one or two reactants. At any time step, a particle that is subject to an unary reaction will react with probability *p*(*λ*; *τ*). If there are two products, they are placed within a sphere of specified radius *R*_u_ around the educt’s position **x**_0_. This is achieved by randomly selecting an orientation **n** ∈ ℝ^3^, distance *d* ≤ *R*_u_, and weights *w*_1_ ≥ 0,*w*_2_ ≥0, s.t. *w*_1_ + *w*_2_ = 1. The products are placed at **x**_1_ = **x**_0_ + *dw*_1_**n** and **x**_2_ = **x**_0_ − *dw*_2_**n**. Per default, *w*_1_ = *w*_2_ = 0.5 and the distances *d* are drawn such that the distribution is uniform with respect to the volume of the sphere. When it is necessary to produce new particles, we suggest to use to define a producing particle A and use the unary reaction A ⇀A + B with corresponding placement weights *w*_1_ = 0*, w*_2_ = 1 so that the A particle stays at its position.

The basic binary reaction scheme is the Doi scheme [14, 46] in which a reactive complex is defined by two reactive particles being in a distance of *R_b_* or less, where *R_b_* is a parameter, e.g., see Fig. 1b Fusion or Enzymatic reaction. The reactive complex then forms with probability *p*(*λ*; *τ*) while the particles are within distance.

Optionally ReaDDy 2 can simulate reversible reactions using the reversible iPRD-DB scheme developed in [21]. This scheme employs a Metropolis-Hastings algorithm that ensures the reversible reaction steps to be made according to thermodynamic equilibrium by accounting for the system’s energy in the educt and product states.

### 2.4 Topologies

Topologies are a way to group particles into superstructures. For example, large-scale molecules can be represented by a set of particles corresponding to molecular domains assembled into a topology. A topology also has a set of potential energy terms such as bond, angle, and torsion terms associated. The specific potential terms are implied by finding all paths of length two, three, and four in the topology connectivity graph. The sequence of particle types associated to these paths then is used to gather the potential term specifics, e.g., force constant, equilibrium length or angle, from a lookup table (Fig. 1a).

Reactions are not only possible between particles, but also between a topology and a particle (Fig. 1b) or two topologies. In order to define such reactions, one can register topology types and then specify the consequences of the reaction on the topology’s connectivity graph. We distinguish between global and local topology reactions.

Global topology reactions are triggered analogously to unary reactions, i.e., they can occur at any time with a fixed rate and probability as given in (5). Any edge in the graph can be removed and added. Moreover, any particle type as well as the topology type can be changed, which may result in significant changes in the potential energy. If the reaction causes the graph to split into two or more components, these components are subsequently treated as separate topologies that inherit the educt’s topology type and therefore also the topology reactions associated with it. Such a reaction is the topology analogue of a particle fission reaction.

A local topology reaction is triggered analogously to binary reactions with probability *p*(*λ*; *τ*) if the distance between two particles is smaller than the reaction radius. At least one of the two particles needs to be part of a topology with a specific type. The product of the reaction is then either yielded by the formation of an edge and/or a change of particle and topology types. In contrast to global reactions only certain changes to particle types and graphs can occur:

- Two topologies can fuse, i.e., an additional edge is introduced between the vertices corresponding to the two particles that triggered the reaction.
- A topology and a free particle can fuse by formation of an edge between the vertex of the topology’s particle and a newly introduced vertex for the free particle.
- Two topologies can react in an enzymatic fashion, i.e., particle types of the triggering particles and topology types can be changed.
- Two topologies and a free particle can react in an enzymatic fashion analogously.

In all of these cases the involved triggering particles’ types and topology types can be changed.

### 2.5 Simulation setup and boundary conditions

Once the potentials, the reactions (Fig. 1a,b), and a temperature *T* have been defined, a corresponding simulation can be set up. A simulation box can be periodic or partially periodic, see Fig. 1c. Periodicity in a certain direction means that with respect to that direction particle wrapping and the minimum image convention are applied. Non-periodic directions require a harmonically repelling wall as given in (4).

In order to define the initial condition, particles and particle complexes are added explicitly by specifying their 3D position and type. A simulation can now be started by providing a time step size *τ* and a number of integration steps.

## 3 Software

ReaDDy 2 is mainly written in C++ and has Python bindings making usage, configuration, and extension easy while still being able to provide high performance. To encourage usage and extension of the software, it is Open Source and licensed under the BSD-3 license. It therefore can not only be used in other Open Source projects without them requiring to have a similar license, but also in a commercial context.

### 3.1 Design

The software consists of three parts. The user-visible toplevel part is the python user interface, see Fig. 2a. It is a language binding of the C++ user interface (Fig. 2b) and has additional convenience functionality. The workflow consists out of three steps:

**Figure 2:**
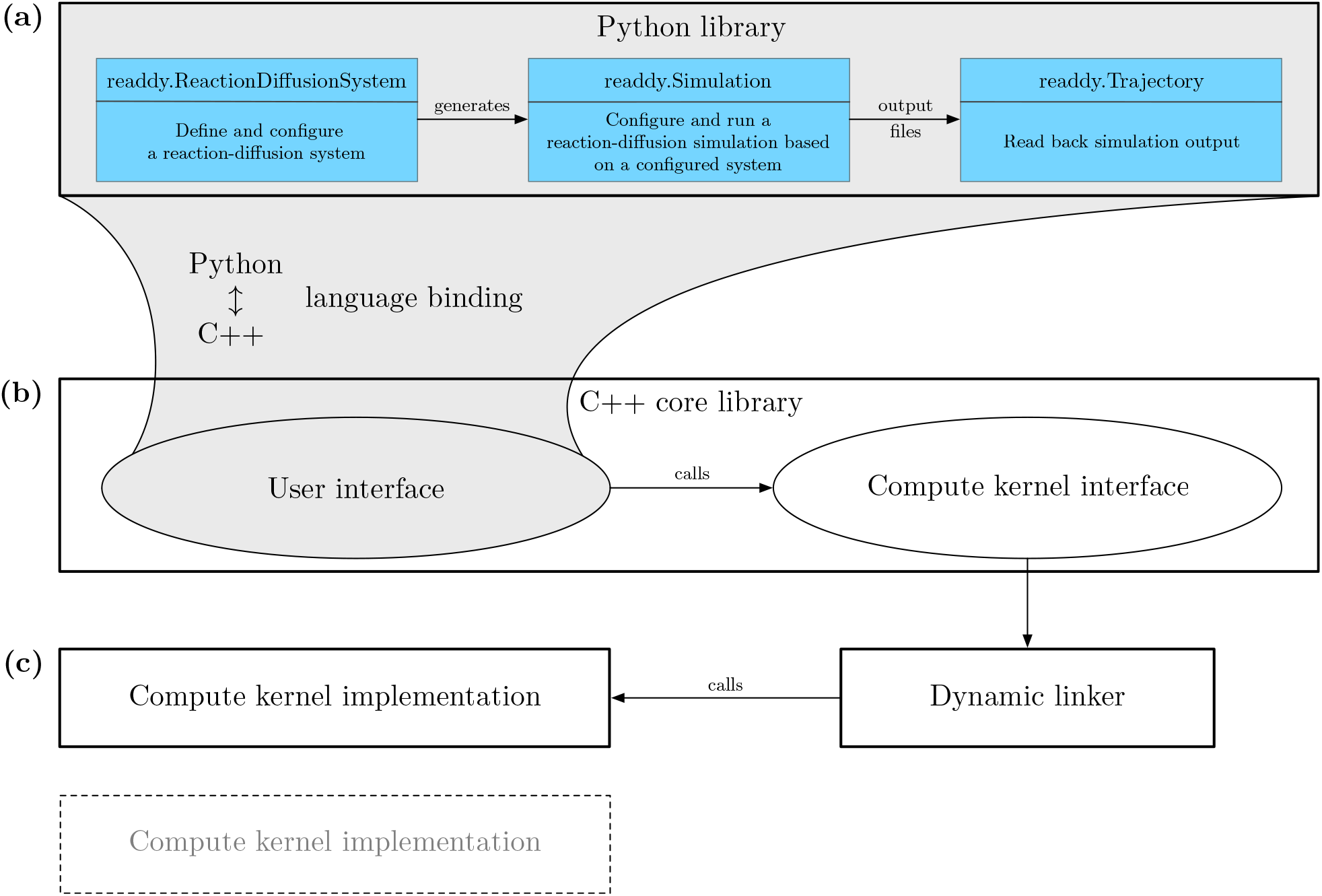
ReaDDy 2 software structure **(a) Python user interface**: Provides a Python binding to the “C++ user interface” with some additional convenience functionality. The user creates a “readdy.ReactionDiffusionSystem” and defines particle species, reactions, and potentials. From a configured system, a “readdy.Simulation” object is generated, which can be used to run a simulation of the system given an initial placement of particles. **(b) C++ core library**: The core library serves as an adapter between the actual implementation of the algorithms in a compute kernel and the user interface. **(c) Compute kernel implementation**: Implements the compute kernel interface and contains the core simulation algorithms. Different compute kernel implementations support different hard- or software environments, such as serial and parallel CPU implementations. The compute kernel is chosen when the “readdy.Simulation” object is generated and then linked dynamically in order to provide optimal implementations for different computing environments under the same user interface.

1. The user is creating a *readdy.ReactionDiffusionSystem*, including information about temperature, simulation box size, periodicity, particle species, reactions, topologies, and physical units. Per default the configurational parameters are interpreted in a unit set well suited for cytosolic environments (lengths in nm, time in ns, and energy in kJ*/*mol), e.g., particles representing proteins in solution. The initial condition, i.e., the positions of particles, is not yet specified.
2. The system can generate one or many instances of *readdy.Simulation*, in which particles and particle complexes can be added at certain positions. When instantiating the simulation object, a compute kernel needs to be selected, in order to specify how the simulation will be run (e.g., single-core or multi-core implementation). Additionally, observables to be monitored during the simulation are registered, e.g., particle positions, forces, or the total energy. A simulation is started by entering a time step size *τ >* 0 in units of time and a number of integration steps that the system should be propagated.
3. When a simulation has been performed, the observables’ outputs have been recorded into a file. The file’s contents can be loaded again into a *readdy.Trajectory* object that can be used to produce trajectories compatible with the VMD molecular viewer [28].

Running a simulation based on the *readdy.Simulation* object invokes a simulation loop. The default simulation loop is given in Alg. 1. Individual steps of the loop can be omitted. This enables the user to, e.g., perform pure PBRD simulations by skipping the calculation of forces. Performing a step in the algorithm leads to a call to the compute kernel interface, see Fig. 2b. Depending on the selected compute kernel the call is then dispatched to the actual implementation. Compute kernel implementations (Fig. 2c) are dynamically loaded at runtime from a plugin directory. This modularity allows ReaDDy 2 to run across many platforms although not every computing kernel may run on a given platform, such as a CUDA-enabled computing kernel. ReaDDy version 2.0.0 includes two iPRD computing kernels: a single threaded default computing kernel, and a dynamically-loaded shared-memory parallel kernel.

The computing kernels contain implementations for the single steps of the simulation loop. Currently, integrator and reaction handler are exchangable by user-written C++ extensions. Hence, there is flexibility considering what is actually performed during one step of the algorithm or even what kind of underlying model is applied.

### 3.2 Performance

To benchmark ReaDDy 2, we use a reactive system with three particle species A, B, and C introduced in [41] with periodic boundaries instead of softly repelling ones. The simulation temperature is set to *T* = 293 K and the diffusion coefficients are given by *D*_A_ = 143.1 *μ*m^2^ s^*−*1^, *D*_B_ = 71.6 *μ*m^2^ s^*−*1^, and *D*_C_ = 68.82 *μ*m^2^ s^*−*1^. Particles of these types are subject to the two reactions A + B ⇀C with microscopic association rate constant *λ*_on_ = 10^*−*3^ ns^*−*1^ and reaction radius *R*_1_ = 4.5 nm, and C ⇀A + B with microscopic dissociation rate constant *λ*_off_ = 5 ×10^*−*5^ ns^*−*1^ and dissociation radius *R*_2_ = *R*_1_. Particles are subject to an harmonic repulsion interaction potential which reads

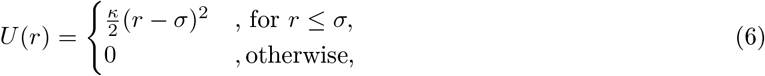

where *σ* is the distance at which particles start to interact and *κ* = 10 kJ mol^*−*1^ nm^*−*2^ is the force constant. The interaction distance *σ* is defined as sum of radii associated to the particles’ types, in this case *r*_A_ = 1.5 nm, *r*_B_ = 3 nm, and *r*_C_ = 3.12 nm. All particles are contained in a cubic box with periodic boundaries. The edge length is chosen such that the initial number density of all particles is *ρ*_tot_ = 3141 nm^*−*3^. This total density is distributed over the species, such that the initial density of A is *ρ*_A_ = *ρ*_tot_*/*4, the initial density of B is *ρ*_B_ = *ρ*_tot_*/*4, and the initial density of C is *ρ*_C_ = *ρ*_tot_*/*2. For the chosen microscopic rates these densities roughly resemble the steady-state of the system. The performance is measured over a simulation timespan of 300 ns which is much shorter than the equilibration time of this system. Thus the overall number of particles does not vary significantly during measurement and we obtain the computation time at constant density.

#### Algorithm 1

ReaDDy 2 default simulation loop. Each of the calls are dispatched to the compute kernel, see Fig. 2. Furthermore, the user can decide to switch off certain calls in the simulation loop while configuring the simulation.

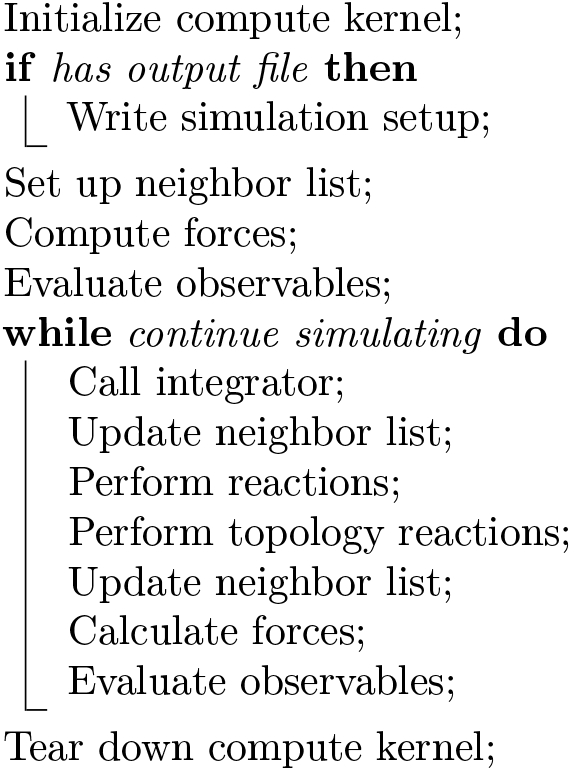

In the following the benchmark results are presented. A comparison between the sequential reference compute kernel, the parallel implementation, and the previous Java-based ReaDDy 1 [42] is made with respect to their performance when varying the number of particles in the system keeping the densityconstant. Since the particle numbers fluctuate the comparison is based on the average computation time per particle and per integration step (Fig. 3). The sequential kernel scales linearly with the number of particles, whereas the parallelized implementation comes with an overhead that depends on the number of threads. The previous Java-based implementation does not scale linearly for large particle numbers, probably owing to Java’s garbage collection. The parallel implementation starts to be more efficient than the sequential kernel given sufficiently many particles.

**Figure 3:**
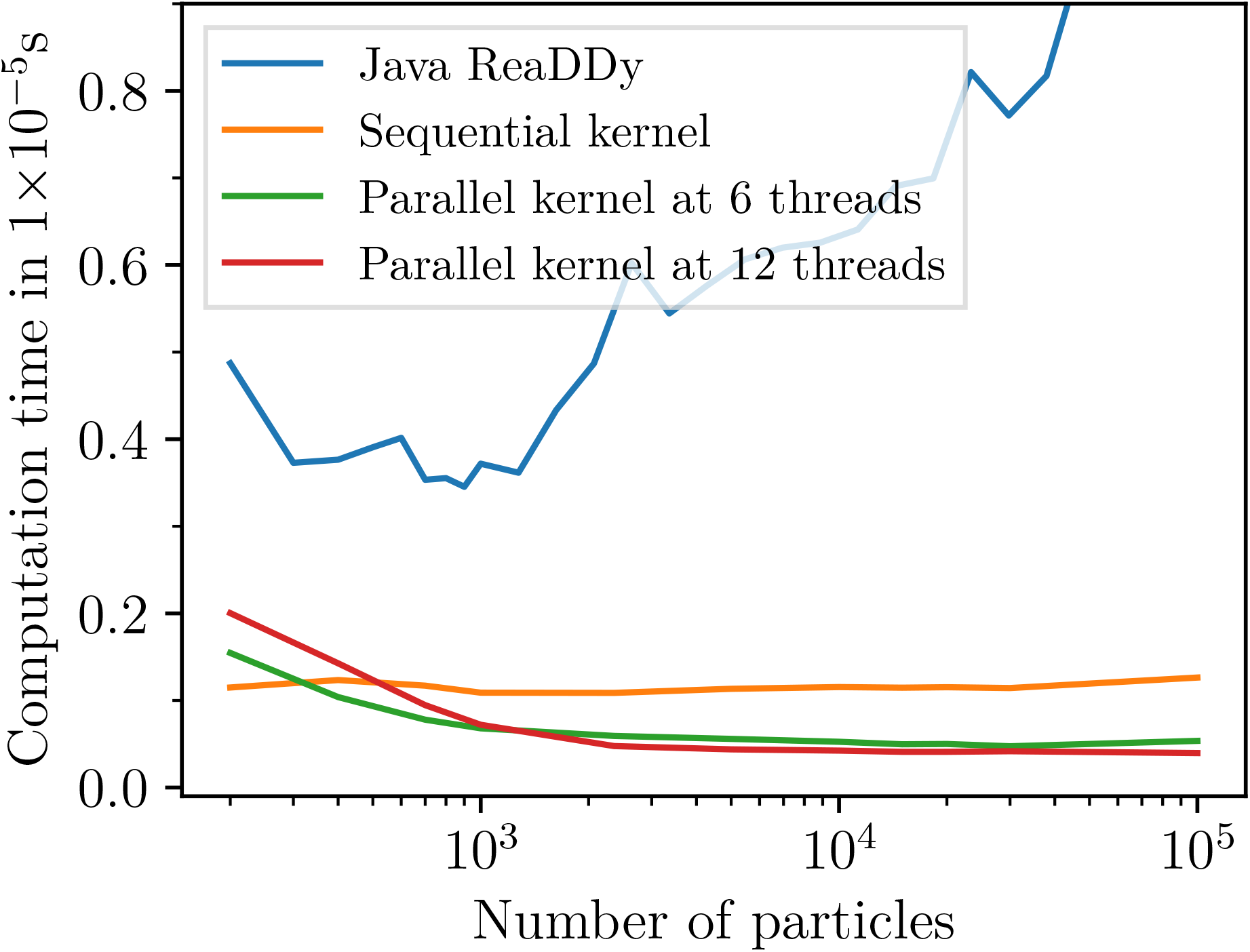
Average computation time per particle and integration step for the benchmark system of Sec. 3.2 using a machine with an Intel Core i7 6850K processor, i.e., six physical cores at 3.8 GHz, and 32 GB DDR4 RAM at 2.4 GHz (dual channel). The number of particles is varied, but the particle density is kept constant. The sequential kernel (orange) has a constant per-particle CPU cost independent of the particle number. For large particle numbers, the parallel kernels are a certain factor faster (see scaling plot Fig. 4). For small particle numbers of a few hundred the sequential kernel is more efficient. ReaDDy 2 is significantly faster and scales much better than the previous Java-based ReaDDy 1 [42].

Fig. 4 shows the strong scaling behavior of the parallel kernel, i.e. the speedup and efficiency for a fixed number of particles as a function of the used number of threads. For sufficiently large particle numbers, the kernel scales linear with the number of physical cores and an efficiency of around 80%. In hyperthreading mode, it then continues to scale linear with the number of virtual cores with an efficiency of about 55–60%.

**Figure 4:**
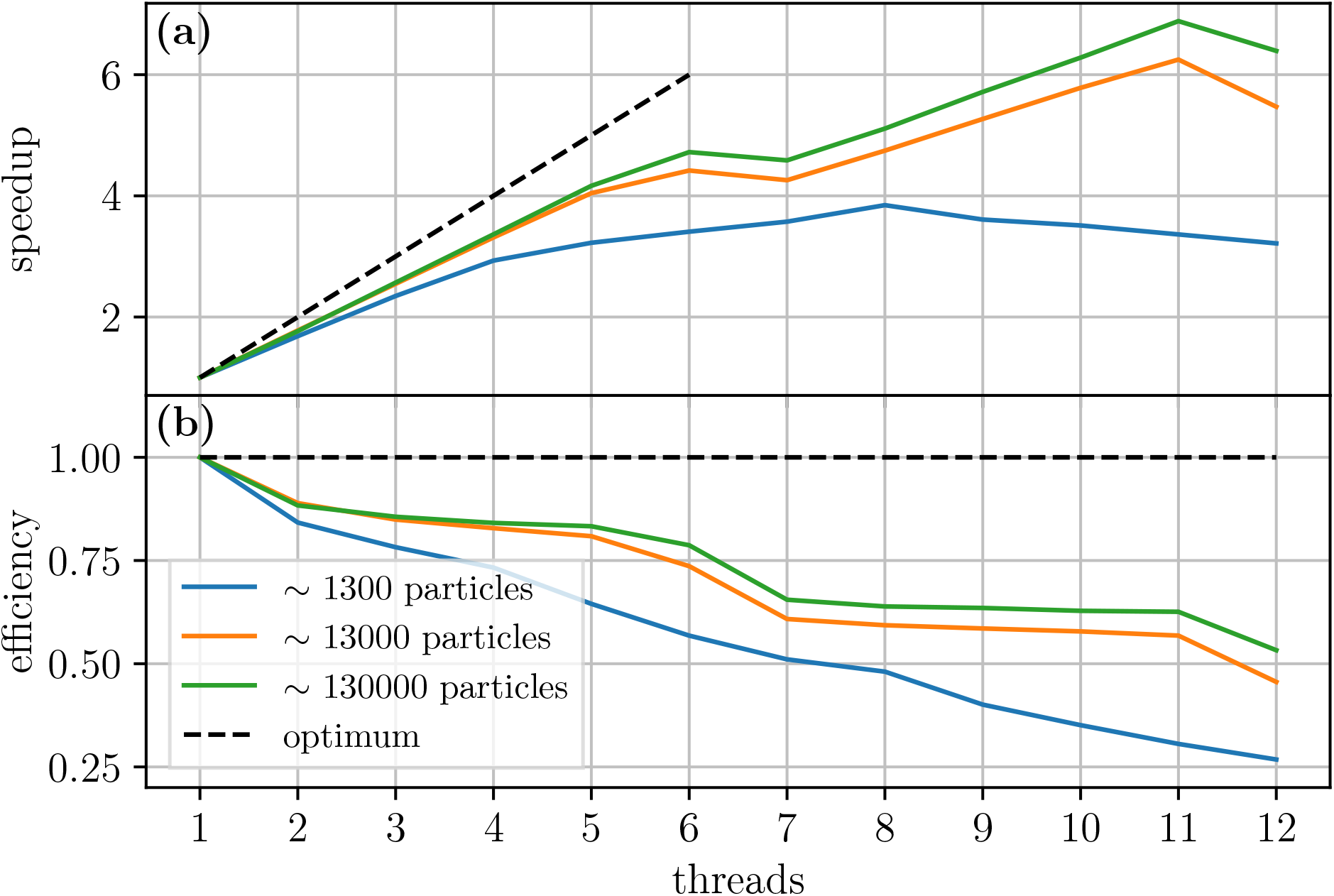
Parallel speedup and efficiency of the benchmark system of Sec. 3.2 as a function of the number of cores using the machine described in Fig. 3. **(a)** Speedup with different numbers of cores compared to one core. Optimally one would like to have a speedup that behaves like the identity (black dashed line). **(b)** Efficiency is the speedup divided by the number of threads, i.e., how efficiently the available cores were used.

The number of steps per day for a selection of particle numbers and kernel implementation is displayed in Tab. 1. For a system with 13, 000 particles and a time step size of *τ* = 1 ns (e.g., membrane proteins [41]), a total of 17 ms simulation time per day can be collected on a six-core machine (Fig. 3 for details). The current ReaDDy kernels are thus suited for the detailed simulation of processes in the millisecond- to second timescale, which include many processes in sensory signalling and signal transduction at cellular membranes.

**Table 1:** Number of time steps per day for benchmark system of Sec. 3.2 using the machine described in Fig. 3. In case of the parallelized implementation the peak performance with respect to the number of threads is shown.

| Approximate<br>number of particles | Steps per day<br>sequential kernel | Peak performance steps per day<br>parallel kernel | Number of<br>threads |
| --- | --- | --- | --- |
| 250 | $2.8 \times 10^8$ | $2.6 \times 10^8$ | 4 |
| 1000 | $7.9 \times 10^7$ | $1.2 \times 10^8$ | 7 |
| 13000 | $5.6 \times 10^6$ | $1.7 \times 10^7$ | 11 |
| 40000 | $1.8 \times 10^6$ | $6.3 \times 10^6$ | 11 |

## 4 Results

In the following, several aspects of the model applied in ReaDDy 2 are validated and demonstrated by considering different application scenarios and comparing the results to analytically obtained results, simulations from other packages, or literature data.

### 4.1 Reaction kinetics and detailed balance

We simulate the time evolution of particle concentrations of the benchmark system described in Sec. 3.2. In contrast to the benchmarks, the considered system initially only contains A and B particles at equal numbers. It then relaxes to its equilibrium mixture of A, B, and C particles (Fig. 5). Since the number of A and B molecules remain equal by construction, only the concentrations of A and C are shown.

**Figure 5.**
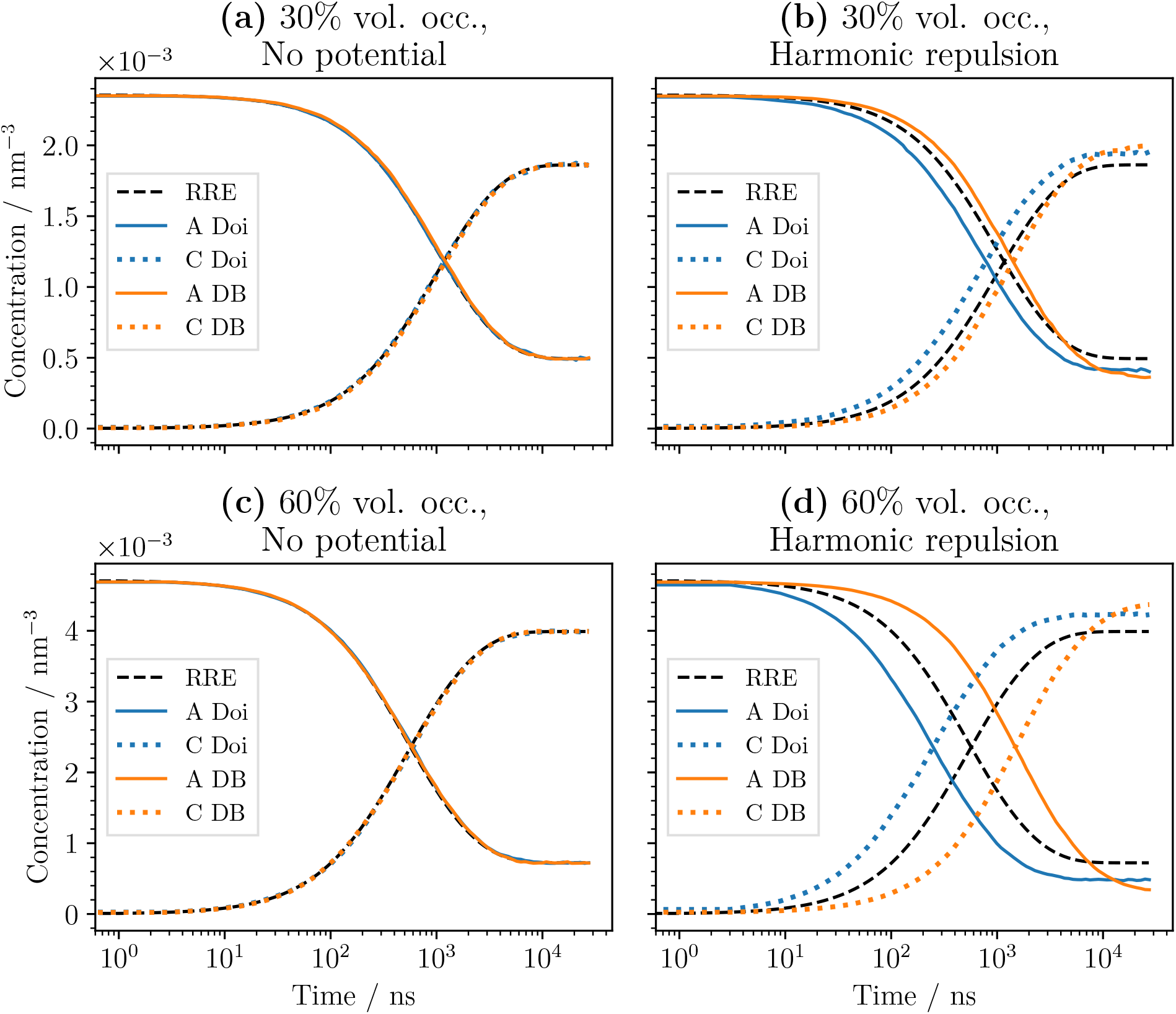
Concentration time series of a the reaction-diffusion system introduced in Sec. 3.2 with the reversible reaction A + B ⇋C. Compared are cases with and without harmonic repulsion (6). Additionally we compare two different reaction mechanisms, the Doi reaction scheme and the detailed balance (iPRD-DB) method for reversible reactions. **(a)** 30% volume occupation and no interaction potentials. **(b)** 30% volume occupation with harmonic repulsion between all particles. **(c)** 60% volume occupation and no interaction potentials. **(d)** 60% volume occupation with harmonic repulsion between all particles.

In addition we compare the solutions with and without harmonic repulsion potentials (6) between all particles, as well as two different methods for executing the reactions: The Doi reaction scheme as described in Sec. 2.3 and the detailed-balance reaction scheme iPRD–DB described in [21].

In contrast to Sec. 3.2, we construct a macroscopic reference system with rate constants *k*_on_ = 3.82 *×* 10^*−*1^ nm^3^s^*−*1^ and *k*_off_ = 5 *×* 10^*−*5^ s^*−*1^ resembling a cellular system. The microscopic reaction rate constants *λ*_on_ and *λ*_off_ are then chosen with respect to the reference system taking interaction potentials between A and B into account by a method presented in [21] under the assumption that the system is sufficiently well-mixed. This yields *λ*_off_ = 5 ×10^*−*5^ ns^*−*1^ for the microsopic dissociation rate constant. The microscopic association rate constant reads *λ*_on_ = 10^*−*3^ ns^*−*1^ for the noninteracting system and *λ*_on_ = 2.89 ×10^*−*3^ ns^*−*1^ for the interacting system. Note that for non-reversible binary reactions without interaction potentials the formula provided by [13, 19] describes the relation between *λ* and *k* for slow diffusion encounter. In the case of non-reversible binary reactions with interaction potentials and slow diffusion encounter such a relation can still be numerically computed [12].

Using the macroscopic rate constants *k*_on_ and *k*_off_, a solution can be calculated for the mass-action reaction rate equations (RRE). This solution serves as a reference for the noninteracting system (no potentials), because the system parameters put the reaction kinetics in the mass-action limit.

In the noninteracting system, the ReaDDy solution and the RRE solution indeed agree (Fig. 5a, c). In the case of interacting particles, see Fig. 5b, d, an exact reference is unknown. We observe deviations from the RRE solution that become more pronounced with increasing particle densities. A difference between the two reaction schemes can also be seen. The Doi reaction scheme shows faster equilibration compared to RRE for increasing density, whereas the iPRD-DB scheme shows slower equilibration, as it has a chance to reject individual reaction events based on the change in potential energy. Thus an increased density leads to more rejected events, consistent with the physical intuition that equilibration in a dense system should be slowed down. Furthermore the equilibrated states differ depending on the reaction scheme, showing a dependence on the particle density. For denser systems the iPRD-DB scheme favors fewer A and B particles than the Doi scheme, consistent with the density-dependent equilibria described in [21].

### 4.2 Diffusion

Next we simulate and validate the diffusive behavior of noninteracting particle systems and the subdiffusive behavior of dense interacting particle system. The simulation box contains the particles with diffusion coefficient *D* and is equipped with softly repelling walls, in order to introduce finite size effects. The observations are carried out without interaction potentials and with harmonic repulsion potentials (6) and, if available, compared to an analytic solution. Particle diameters are *σ*, time *t* is given in units of *σ*^2^*/D*, length *x* is given in units of *σ*, and energy is given in units of *k_B_T*. The cubic box has an edge length of *ℓ≈* 20.6*σ*.

The noninteracting particle simulation has a mean square displacement of particles in agreement with the analytic solution given by Fick’s law for diffusion in three dimensions

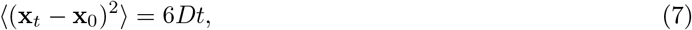

where **x** is the position of a particle and *t* is time (Fig. 6). For long timescales *t* ≥10^1^ transport is obstructed by walls, resulting in finite size saturation.

**Figure 6.**
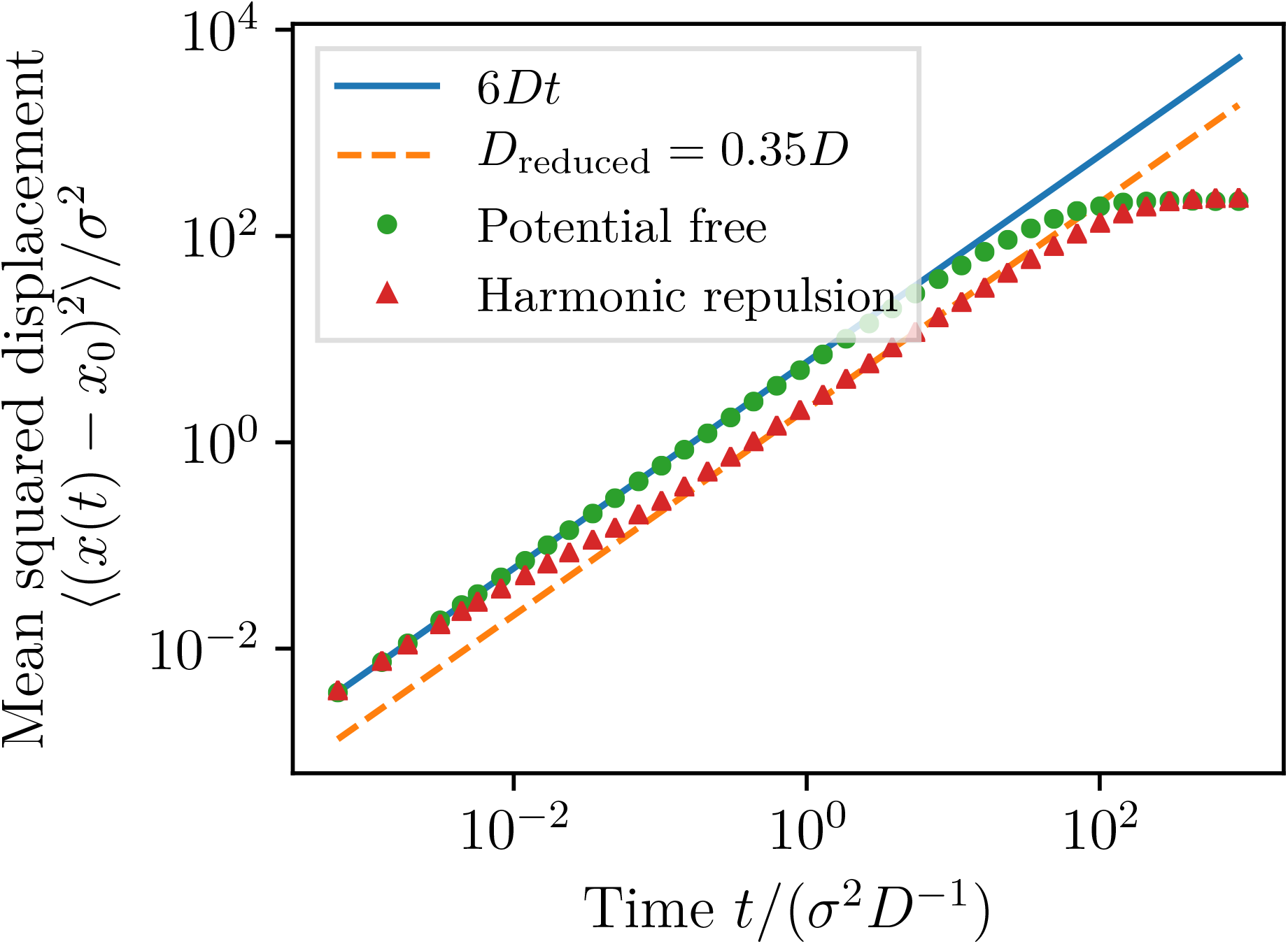
Mean squared displacement as a function of time for multiple particles diffusing with diffusion coefficient *D* in a cubic box with harmonically repulsing walls. Without interaction potentials, the mean square displacement agrees with the analytical result for free diffusion, see Eq. (7). Using harmonic repulsion potentials between all particles results in reduced mobility. On timescales *t* ∈ [10^*−*1^, 10^1^) the mean squared displacement is linear with a reduced effective diffusion coefficient, indicated by the dashed line.

Fig. 6 also shows that more complex transport can be modeled, as, e.g., found in crowded systems. Particles interact via harmonic repulsion (Eq. 6) with a force constant of *κ ≈*79*k_B_Tσ^−^*^2^, *σ* being the particle diameter. The particle system is dense, having a volume occupation of 84%. In such a dense particle system, the mean squared displacement differs significantly from the analytical result for free diffusion after an initial time *t* ≥10^*−*2^ in which particles travel their mean free path length with diffusion constant *D*. At intermediate timescales *t* ∈ [10^*−*2^, 10^*−*1^), particle transport is subdiffusive due to crowding. At long timescales, *t* ∈ [10^*−*1^, 10^1^), the particles are again diffusive with an effective diffusion coefficient that is reduced to reflect the effective mobility in the crowded systems. For large timescales *t* ≥10^1^ finite size saturation can be observed again, as previously.

### 4.3 Thermodynamic equilibrium properties

We validate that ReaDDy 2’s integration of equations of motion yields the correct thermodynamics of a Lennard-Jones colloidal fluid in an (*N, V, T*) ensemble. To this end, we simulate a system of *N* particles confined to a periodic box with volume *V* at temperature *T*. The results and comparisons with other simulation frameworks and analytical results are shown in Tab. 2. The particles interact via the Lennard-Jones potential

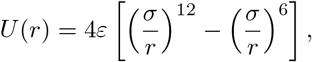

with *ε* being the depth of the potential well and *σ* the diameter of particles. The potential is cut off at *r_C_* = 4*σ* and shifted to avoid a discontinuity. The rescaled temperature is *T ^*^* = *k_B_T* ϵ^*−*1^ = 3. We perform simulations of the equilibrated Lennard-Jones system for 10^6^ integration steps with rescaled time step size *τ ^*^* = 10^*−*4^. Time units are *σ*^2^*/D* and are determined by the self-diffusion coefficient *D* of the particles. We measure the rescaled pressure *P ^*^*= *Pσ*^3^*ε^−^*^1^ by estimating the virial term from forces acting in the system as described in [3]. Additionally we measure the rescaled potential energy per particle *u^*^* = *UN ^−^*^1^*ε^−^*^1^. Both pressure and potential energy are calculated every 100th time step. This sampling gives rise to the mean and its error of the mean given for the ReaDDy 2 results in Tab. 2. Comparing HALMD [11] and ReaDDy 2, the latter shows larger energy and pressure in the third decimal place for the lower density *ρ^*^* = 0.3. For the higher density *ρ^*^* = 0.6 pressure differs in the first decimal place and energy in the second. This can be explained by ReaDDy 2 using an Euler scheme (3) to integrate motion of particles, which has a discretization error of first order in the time step size 𝓞 (*τ*). On the other hand HALMD uses a Velocity-Verlet method [44], which has a discretization error of second order in the time step size 𝓞(*τ* ^2^).

**Table 2:** Thermodynamic equilibrium properties of a Lennard–Jones colloidal fluid in a (*N, V, T*) ensemble. Results of the ReaDDy 2 framework are compared to other simulation frameworks and analytical results for validation.

| | density $\rho^*$ | pressure $P^*$ | energy $u^*$ |
| --- | --- | --- | --- |
| ReaDDy 2 | 0.3 | $1.0253 \pm 0.0004$ | $-1.6704 \pm 0.0003$ |
| HALMD [11] | 0.3 | $1.0234 \pm 0.0003$ | $-1.6731 \pm 0.0004$ |
| Johnson et al. [29] | 0.3 | $1.023 \pm 0.002$ | $-1.673 \pm 0.002$ |
| Ayadim et al. [7] | 0.3 | 1.0245 | -1.6717 |
| ReaDDy 2 | 0.6 | $3.711 \pm 0.002$ | $-3.2043 \pm 0.0004$ |
| HALMD [11] | 0.6 | $3.6976 \pm 0.0008$ | $-3.2121 \pm 0.0002$ |
| Johnson et al. [29] | 0.6 | $3.69 \pm 0.01$ | $-3.212 \pm 0.003$ |
| Ayadim et al. [7] | 0.6 | 3.7165 | -3.2065 |

### 4.4 Topology reactions

Finally, we illustrate ReaDDy 2’s ability to model complex reactions between multi-particle complexes, called “topology reactions”. We model polymers as linear chains of beads, held together by harmonic bonds and stiffened by harmonic angle potentials. These beads can have two different particle types. Either they are head particles and located at the ends of a polymer chain or they are core particles and located between the head particles, as shown in Fig. 7a, c in blue and orange, respectively.

**Figure 7.**
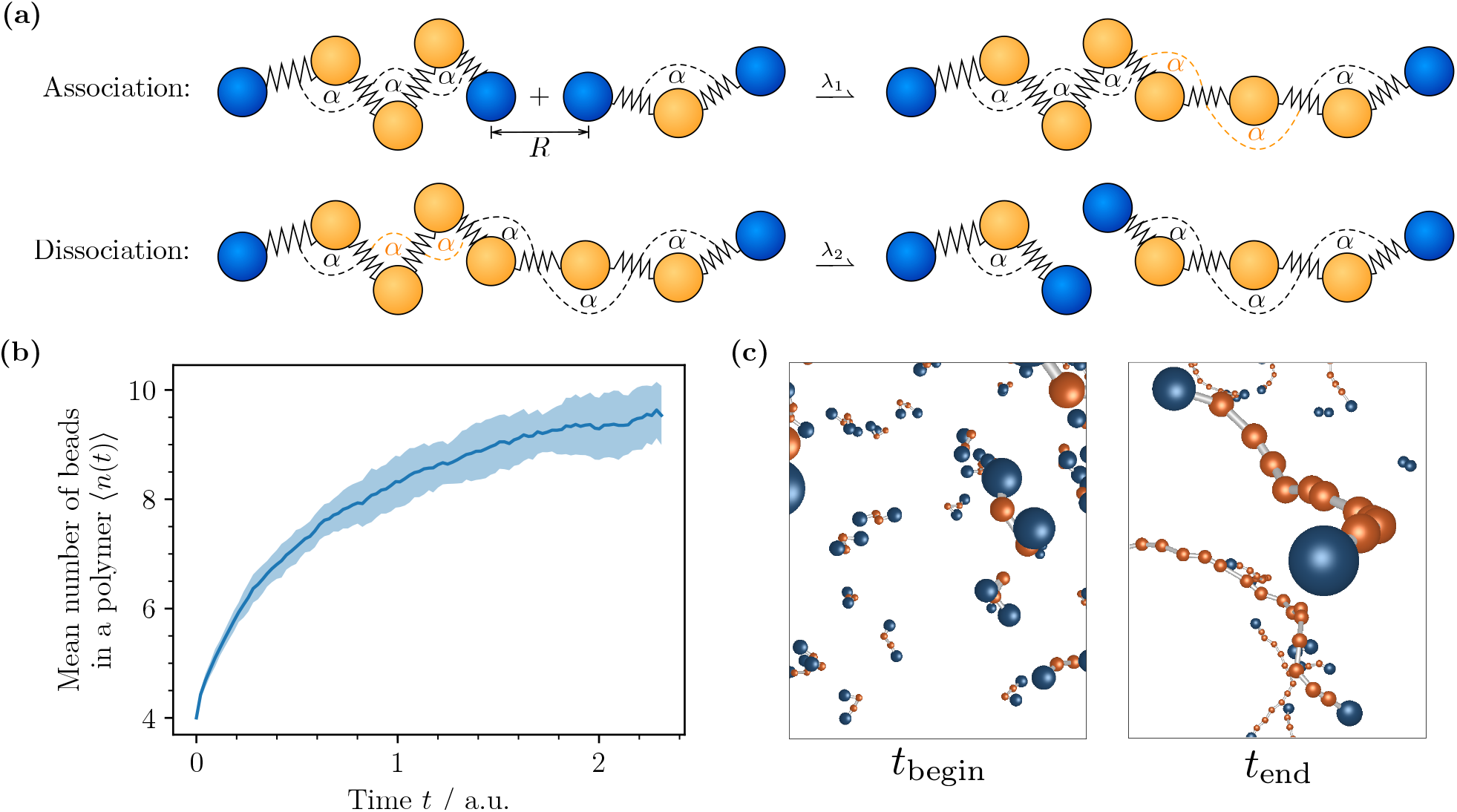
Illustrative simulation of polymer assembly/disassembly using topology reactions. **(a)** Sketch of the involved topology reactions. *Association*: When two ends of different topologies come closer than *R*, there is a rate *λ*_1_ that an edge is formed. *Dissociation*: The inverse of association with a rate *λ*_2_ and a randomly drawn edge that is removed. **(b)** The number of beads in a polymer 〈*n*(*t*) 〉 over time averaged over 15 realizations. **(c)** Two representative particle configurations showing the initial state and the end state at time *t*_begin_and *t*_end_, respectively.

There are two different topology reactions in the system (Fig. 7a):

1. Association: Two nearby head particles (distance ≤*R*) can connect with rate *λ*_1_. The topology is changed by adding an edge between the connected particles, resulting in the addition of one bond and two angle potentials. Additionally, the particle types of the two connected particles change from “head” to “core”.
2. Dissociation: A chain with *n* particles can dissociate with microscopic rate *nλ*_2_, such that longer chains have a higher probability to dissociate than shorter chains. When a dissociation occurs, a random edge between two core particles is removed. The particle types of the respective core particles are changed to “head”. As a result, the graph decays into two connected components which subsequently are treated as autonomous topology instances.

The temporal evolution of the average length of polymer chains is depicted in Fig. 7b. The simulation was performed 15 times with an initial configuration of 500 polymers containing four beads each. After sufficient time ⟨*n*(*t*) ⟩ reaches an equilibrium value. Over the course of the simulation the polymers diffuse and form longer polymers. This can also be observed from the two snapshots shown in Fig. 7c, depicting a representative initial configuration at *t*_begin_ and a representative configuration at the end of the simulation at time. In that case, there are polymers of many different lengths.

## 5 Conclusions and outlook

We have described the iPRD simulation framework ReaDDy 2, a simulation framework for combined particle interaction dynamics and reaction kinetics, which permits to conduct highly realistic simulations of signal transduction in crowded cellular environments or chemical nanoreactors with complex geometries. ReaDDy 2 follows up upon and significantly extends the simulation package ReaDDy 1. ReaDDy 2 is significantly faster than its predecessor, it can be easily installed as a Python conda package, and it can be flexibly used and reconfigured via its Python interface.

We have conducted a set of numerical studies, showing that ReaDDy 2 produces quantitatively accurate results where references from analytical solutions or other simulation packages were available, and physically meaningful results where reference solutions were not available.

For a quick and easy start into simulating and developing with ReaDDy 2 step by step tutorials, sample code, and further details are available online (https://readdy.github.io/). The software itself is Open Source and available under a permissive licence in order to enable a broad group of people to run simulations without forcing them to make their own work public.

ReaDDy 2 has been designed to be easily extensible. Planned extensions include simulation kernels for specialized hardware platforms, such as graphics processors and highly parallel HPC environments. Also planned is a MD-GFRD integrator [38] to speed up computations in dilute systems, and a particle-based membrane model as described in [36] that reproduces mechanical properties of cellular membranes.

## Acknowledgements

The authors are grateful to the Center for Theoretical Biological Physics (CTBP, supported by NSF PHY-1427654) at Rice University for hosting their sabbatical visit, during which part of this work was performed.

We gratefully acknowledge funding from Deutsche Forschungsgemeinschaft (SFB 958 / Project A04, TRR 186 / Project A12, SFB 1114 / Project C03), Einstein Foundation Berlin (ECMath Project CH17) and European Research Council (ERC CoG 772230 “ScaleCell”). We are grateful for inspiring discussions with Manuel Dibak, Luigi Sbailò, Mohsen Sadeghi, Felix Höfling and Christof Schütte.

